# Inferring norepinephrine dynamics from partial observations reveals the temporal structure of elevations during arousal

**DOI:** 10.64898/2026.03.29.715097

**Authors:** Erin Neyhart, Brandon R. Munn, Na Zhou, Peilin Yang, Jiesi Feng, Yulong Li, James M. Shine, Jacob Reimer

## Abstract

Hemodynamic artifacts present a significant challenge for two-photon fluorescence imaging of genetically encoded reporters, particularly when the timescale of relevant measurements matches those of vascular dynamics. This is an acute challenge for sensors in which the hemodynamic artifact is of comparable magnitude to the biological signal of interest. However, standard correction methods, such as isobestic recording or repeated experiments, are often impractical. Here we introduce a tiered framework for inferring norepinephrine (NE) dynamics across varying levels of recording information. First, we verify that dual-channel recording using an inert fluorescent reporter alongside the neuromodulator indicator enables direct hemodynamic correction within the same recording session. For contexts in which a dedicated reference channel is unavailable, we trained an LSTM-based model that predicts and removes hemodynamic contributions post-hoc from the recorded NE signal and behavioral variables. Finally, we show that key features of NE dynamics can be recovered from behavioral variables alone, providing an estimate of neuromodulatory state even when fluorescence recordings are unavailable. These methods enabled simultaneous multi-spectral measurements of axonal activity and neuromodulator release via simultaneous two-photon imaging of LC noradrenergic axons and extracellular NE in the same field of view. Cortical NE signals are graded with respect to behavioral intensity, scaling with both locomotion duration and pupil dilation amplitude. As expected, axonal activity precedes increases in ambient NE levels, but NE peaks later within a run and remains elevated after axonal activity has subsided, suggesting that extracellular NE integrates LC output over time rather than tracking instantaneous LC firing. Together, these findings demonstrate that accurate hemodynamic correction is essential for interpreting NE dynamics, and reveal a clearer view of the temporal structure of cortical norepinephrine signaling.

## INTRODUCTION

The locus coeruleus (LC) is the major supplier of norepinephrine (NE) to the cerebral cortex, and the LC-noradrenergic system has long been associated with states of arousal, attention, and behavioral flexibility^1,2^.Through widespread LC projections across the brain^3^, NE acts as a powerful neuromodulator, dynamically shaping neural excitability^4,5^ and coordinating large-scale network activity in response to both internal and external demands^6–8^. Until recently, directly measuring extracellular NE dynamics with high spatiotemporal resolution has been difficult, with studies relying on cyclic voltammetry (which has trouble differentiating catecholamines), slow methods such as microdialysis, or indirect proxies, such as LC axon imaging with calcium indicators like GCaMP. The development of genetically encoded fluorescent NE sensors such as GRAB_NE_ ^9^ and nLight^10^ has largely overcome this barrier, enabling direct, real-time monitoring of extracellular NE *in vivo* and spurring growing interest in the spatiotemporal structure of NE signals across the cortex and their relationship to behavior and brain state transitions^11–14^.

An ongoing challenge in interpreting bulk fluorescence signals from *in vivo* two-photon imaging is contamination by hemodynamic artifacts, including changes in blood volume, oxygenation, and vessel diameter that modulate light absorption and scattering, thereby distorting the detected fluorescence signal. These effects are particularly problematic in the context of neuromodulator sensors, because hemodynamics are tightly coupled to the same behavioral and brain state variables^15^ that drive neuromodulatory activity, as well as NE release itself^16,17^.

While hemodynamic confounds and the need to correct for them have long been recognized in widefield fluorescence imaging and fiber photometry^18–21^, they are not always explicitly accounted for in two-photon recordings of bulk fluorescence indicators. Indeed, many studies employing genetically encoded NE sensors have not applied explicit hemodynamic corrections^12–14,22^. Importantly, recent work has demonstrated that hemodynamic occlusion effects can be substantial, with magnitudes sometimes comparable to those of activity-dependent fluorescence signals and exhibiting spatial heterogeneity across cortical depth and regions^17,23,24^. The issue is compounded for sensors in which the hemodynamic artifact is of comparable magnitude to the biological signal of interest: a regime in which uncorrected recordings may be dominated by artifactual variance.

Solutions to hemodynamic contamination in two-photon imaging include isobestic illumination (using a second light source), or characterizing the hemodynamic contribution through separate control experiments. Both approaches carry significant practical limitations: isobestic illumination requires a second tunable two-photon light source, and separate control experiments would ideally need to be repeated across all relevant imaging conditions given that hemodynamic occlusion varies across cortical regions and depths. Here we validate another established solution which has been used routinely in other contexts: simultaneous two-channel recording with an inert fluorescent reporter, which provides a real-time hemodynamic reference within the same experiment. This approach cannot be applied universally, as many recording scenarios require multi-color recordings of other dynamic signals. To address this, we used a large set of dual-color recordings with GRAB_NE_ and an inert sensor to train a neural network that infers and removes hemodynamic artifacts when only the GRAB_NE_ signal, pupil, and treadmill activity is known. This approach enables correction of datasets lacking a static control channel. Finally, in the most information-limited regime, we show that key features of NE dynamics can be recovered from behavioral variables alone, providing a useful estimate of neuromodulatory state even when fluorescence recordings are unavailable.

Applying these methods, we uncover previously unrecognized features of the temporal organization of cortical NE dynamics, including systematic scaling of NE magnitude with behavior, and persistent NE elevations that outlast LC axon activity. These findings highlight how inaccuracies in hemodynamic correction can obscure fundamental aspects of neuromodulatory signaling, and underscore the importance of accurate signal recovery for interpreting NE function across scales. By making robust NE measurement accessible across a broader range of experimental contexts, this framework expands the utility of the growing body of genetically encoded fluorescent NE datasets and facilitates more reliable comparisons across studies, brain regions, and behavioral conditions.

## RESULTS

### Validation of two-channel correction method

To begin investigating the spatiotemporal dynamics of extracellular cortical NE in relation to arousal states, we performed two-photon recordings with GRAB_NE_ in awake, behaving mice during spontaneous locomotion while simultaneously monitoring pupil size and treadmill speed. Initial recordings in primary visual cortex revealed an unexpected result: rather than an increase in GRAB_NE_ fluorescence reflecting the anticipated increase in NE activity at run onset^12,14,25–27^, we instead observed a decrease in fluorescence, followed by an increase at run offset (Supplemental Figure 1A). Markedly, the magnitude of this run-aligned decreased varied with imaging depth (Supplemental Figure 1B) ^23^. A similar unexpected pattern was observed when aligning signals to spontaneous pupil dilation events outside of running (i.e., during quiescence): We observed a small increase, then larger decrease in NE levels proceeding pupil dilation, in contrast to substantial evidence that NE increases on average just prior to arousal-related pupil dilation^12,14,25–32^.

To determine whether these observed dynamics reflected true NE activity or an imaging artifact, we imaged three spectrally-distinct static fluorophores under identical conditions: GFP, mApple, and GRAB_NE_-mut (“NE-mut”)^22^, a version of GRAB_NE_ with mutations in the receptor domain that abolish NE sensitivity. All three reporters exhibited qualitatively similar fluorescence changes at run onset and dilation, confirming that these dynamics were hemodynamic in origin rather than reflecting true NE activity (Supplemental Figure 1C). Notably, the artifact was more pronounced in GFP and NE-mut, which share the same fluorophore, than in mApple, verifying that the hemodynamic occlusion effect is wavelength-dependent. Critically, the magnitude of the hemodynamic artifact in the GRAB_NE_ signal was comparable to or larger than the anticipated biological signal, making uncorrected recordings difficult or impossible to interpret. This is not a universal feature of genetically encoded neuromodulator sensors: recordings with genetically encoded acetylcholine (ACh) indicators under the same conditions reveal a hemodynamic artifact that is small relative to the biological signal, such that run- aligned elevations are clearly visible even in uncorrected traces^23^ (Supplemental Figure 1D).

While hemodynamic artifacts in fiber photometry are commonly addressed using isobestic correction, in which a second excitation wavelength is used to isolate a hemodynamic reference signal (Figure 1A, top), this approach was not compatible with our setup, which only utilizes a single excitation wavelength. We therefore developed a two-channel recording strategy in which rGRAB_NE_ (“rNE”) and NE-mut were co-expressed in the same cortical volume and imaged simultaneously in separate spectral channels (Figure 1A, bottom and Figure 1B). We also evaluated two alternative approaches: co-expression of GRAB_NE_ with mApple as an inert red-channel reporter (Supplemental Figure 1E), and a two-FOV strategy in which GRAB_NE_ and NE-mut were injected in spatially segregated regions and imaged in separate fields of view simultaneously (Supplemental Figure 1F). However, mApple’s reduced sensitivity to the hemodynamic artifact resulted in systematic undercorrection of the NE signal, and the two-FOV approach was further complicated both by differences in vasculature across the two sites and by the comparatively larger hemodynamic artifact observed in the green GRAB_NE_ sensor. We therefore adopted the rNE and NE-mut two-channel approach as the most promising strategy.

**Figure 1:**
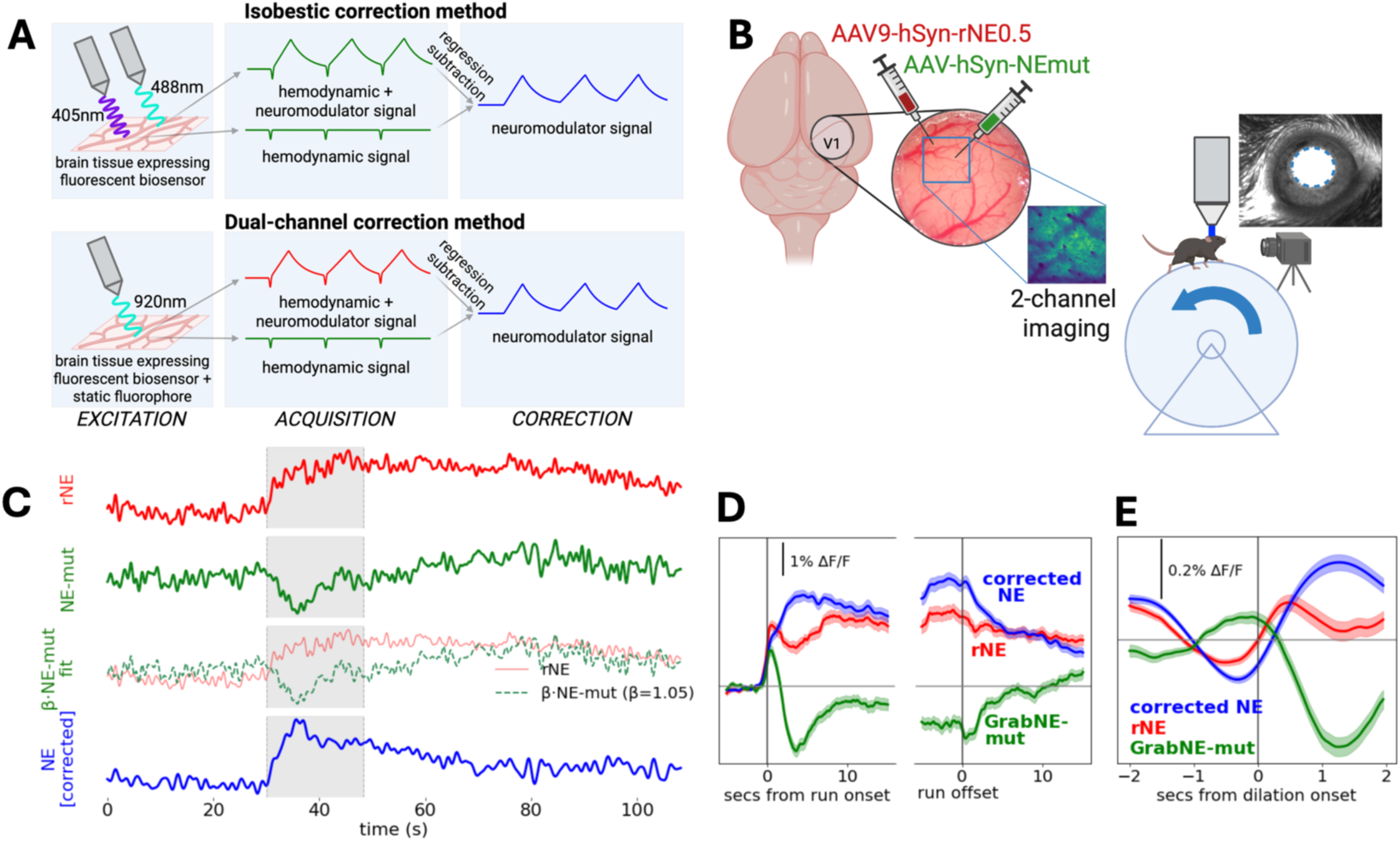
A static fluorophore channel enables direct hemodynamic correction of NE signals. **A)** Comparison of two hemodynamic correction strategies. The isobestic method (top) requires excitation at two wavelengths to isolate a hemodynamic reference, whereas the dual-channel method (bottom) uses a single excitation wavelength and a co-expressed sensor-null construct (NE-mut) as a direct hemodynamic reference. In both cases, regression subtraction of the hemodynamic component from the composite signal yields an estimate of the neuromodulator-driven fluorescence. **B)** Experimental configuration for dual-channel two-photon imaging. rNE and NE-mut are co-injected into mouse cortex through a cranial window and imaged simultaneously. The animal is head-fixed on a treadmill where it is free to run at will, and a separate camera monitors pupil diameter as a proxy for arousal state. **C)** Illustration of the hemodynamic correction algorithm applied to example data. The raw rNE trace (red) contains prominent hemodynamic artifacts that co-vary with the hemodynamic signal represented by the NE-mut (green). A scaling coefficient β is estimated by linear regression between the two channels during stationary periods (locomotion bouts ± buffer excluded; grey shading), then applied to subtract the mean-shifted hemodynamic component from the raw signal. The corrected trace (blue) retains expected neuromodulator dynamics while suppressing apparent vascular artifacts. **D)** Mean run onset- and offset-triggered corrected NE, rNE, and NE-mut activity (n = 422 bouts, 31 ROIs, 6 animals; VIS, SS, and MO cortex). Uncorrected rNE (red) and NE-mut (green) show prominent locomotion-locked hemodynamic transients; corrected NE (blue) isolates the underlying neuromodulatory signal. Shaded areas indicate ± SEM. **E)** Mean dilation-triggered corrected NE, rNE, and NE-mut activity outside of running periods from the same recordings (n = 3,787 dilations, 27 ROIs, 5 animals). Shaded areas indicate ± SEM.

We then adapted a regression-based correction approach^18^, originally developed for single-wavelength reflectance data in widefield imaging, to this dual-channel fluorescence context, regressing the NE-mut signal against the rNE trace and subtracting the scaled hemodynamic component to yield a corrected NE signal (Figure 1C; see Methods). Following correction, the expected increase in NE activity at run onset and return to baseline at run offset were recovered (Fig. 1D), as was the anticipated increase in NE during pupil dilation (Figure 1E). This supports the static fluorophore regression-based correction approach as an effective strategy for isolating true NE dynamics from hemodynamic contamination. We note that all of the proposed methods are likely to have some limitations, and in particular the choice of method may alter the absolute amplitude of the resulting corrected fluorescence signal. Therefore, it is best to limit comparisons to within-session conditions, and direct comparisons of NE magnitude across different correction strategies should be interpreted with caution.

### Applying two-channel correction to characterize behaviorally-linked NE dynamics

Having validated the two-channel correction method in visual cortex, we next asked whether the approach generalized across cortical regions and whether it revealed known or novel features of NE dynamics during behavior. We co-expressed rNE and NE-mut in visual (VIS), somatosensory (SS), or motor (MO) cortex and recorded these signals during spontaneous behavior, and used the NE-mut signal as a hemodynamic reference for correction. In all three regions, the corrected NE signal showed the expected increase at run onset and return to baseline following run offset (Figure 2A), confirming that this correction method generalizes across cortical areas.

**Figure 2.**
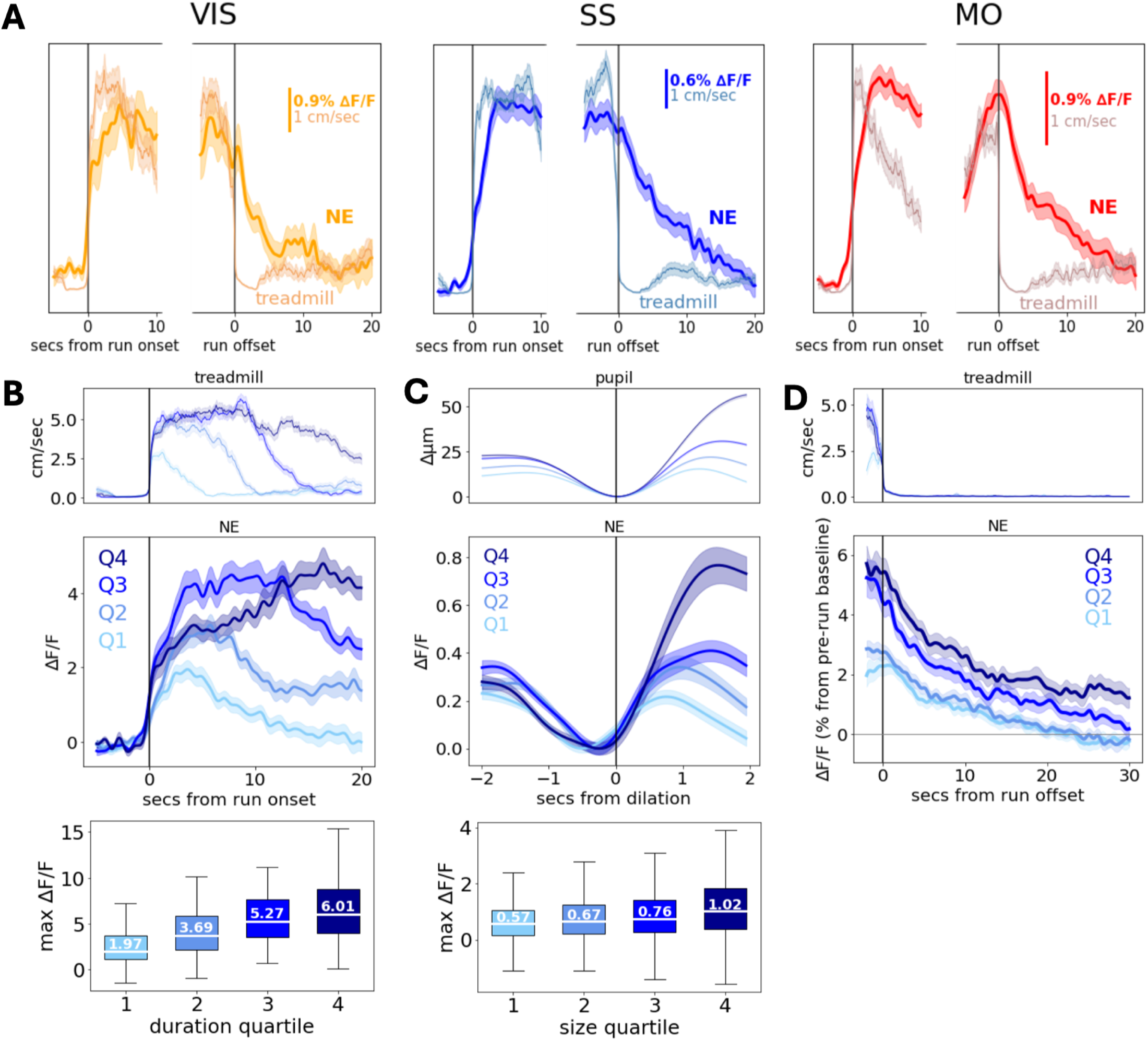
Dual-channel correction reveals graded arousal-coupled NE dynamics. **A)** Mean run onset- and offset- triggered corrected rNE traces for VIS (n=115 run periods from 8 ROIs and 3 mice), SS (n=144 run periods from 10 ROIs and 2 mice), and MO (n=165 run periods from 13 ROIs and 3 mice). **B)** (top) Mean run onset-triggered corrected rNE binned by run duration (n=105-106 periods in each bin) (recordings from all brain areas combined). (bottom) Box plot of peak NE ΔF/F per run period in each bin; median values noted in plot. All pairwise combinations of medians were statistically significant (Kruskal-Wallis test, Bonferroni corrected) except Q3 vs Q4 (1 vs 2 p=2.2×10-6, 1 vs 3 p=1.9×10-14, 1 vs 4 p=6.3×10-18, 2 vs 3 p=0.003, 2 vs 4 p=4.8×10-6, 3 vs 4 p=0.694). **C)** (top) Mean dilation-triggered corrected NE traces outside of running periods sorted by dilation magnitude (n=946-947 dilations in each bin; all brain areas combined). (bottom) Box plot of peak NE ΔF/F per dilation period in each bin; median values noted in plot. All pairwise combinations of medians were statistically significant (Kruskal-Wallis test, Bonferroni corrected) except Q1 vs Q2 (1 vs 2 p=0.062, 1 vs 3 p=1.1×10-6, 1 vs 4 p=7.0×10-25, 2 vs 3 p=0.062, 2 vs 4 p=2.0×10-14, 3 vs 4 p=2.2×10-7). **D)** Mean run offset-triggered corrected NE binned by run duration (n=56-57 periods in each bin; all brain areas combined). Shaded regions throughout indicate ± SEM. Quartile bin definitions for run duration and dilation magnitude are provided in Supplemental Fig. 3.

We next leveraged this dataset to ask whether NE dynamics scale with the intensity of the behavioral event. Stratifying runs across all recorded brain regions by duration into quartiles, we found that longer runs were accompanied by progressively larger peak NE responses (Figure 2B). This relationship was statistically significant across most pairwise quartile comparisons (Mann-Whitney test, Bonferroni corrected) with the exception of Q3 vs Q4, suggesting that while NE magnitude broadly scales with run duration, the relationship may saturate at longer durations. Importantly, this scaling was present in the uncorrected rNE signal and did not appear to be driven by hemodynamics: while the NE-mut signal did show some run duration-dependent structure, with traces stabilizing at different baseline levels across quartiles following an initial artifactual dip at run onset, this pattern is distinct from the progressive peak scaling observed in rNE (Supplemental Figure 2A). We observed a similar scaling relationship for spontaneous pupil dilation events during quiescence (i.e. outside of running periods): stratifying dilations by amplitude revealed that larger dilations were accompanied by larger peak NE responses (Figure 2C), with all quartile pairs significantly different except Q1 vs Q2 and Q2 vs Q3. NE-mut showed no scaling with pupil dilation amplitude (Supplemental Figure 2B), consistent with the locomotion result. Finally, NE responses scaled with run duration not only in peak magnitude but also in their post-offset time course, with longer runs producing more sustained NE elevations (Figure 2D). Together, these results indicate that extracellular NE dynamics are graded with respect to behavioral intensity across both locomotion and pupil-linked arousal, suggesting that NE encodes the magnitude of ongoing behavioral state transitions rather than acting as a simple binary signal.

### Development and validation of an LSTM-based hemodynamic prediction model for post-hoc NE signal correction

While the two-channel correction method described above overcomes the need for a second excitation wavelength and separate control experiments, it still requires simultaneous recording of both the NE-sensitive and static fluorophore channels, which can be a significant constraint for experiments that require multi-color measurements of NE with another dynamic fluorophore. To address this, we developed a model that predicts the hemodynamic component of the fluorescence signal directly from the recorded NE signal and behavioral variables, enabling post-hoc correction without a static control channel.

Consistent with prior work demonstrating the need for computational approaches to separate hemodynamic contributions from fluorescence signals^20^, we trained a long short-term memory (LSTM) network to predict the hemodynamic signal (“NE-mut predicted”) from combinations of four inputs: the raw recorded NE signal (rNE), a spatially-binned version of the NE signal averaged across intensity-defined pixel zones (see Supplemental Figure 4), locomotion, and pupil size behavioral variables (Figure 3A). Leaving out different variables yielded four model variants of increasing complexity. Across all variants, the model achieved moderate but consistent predictive accuracy on held-out test data (Figure 3B). The full model incorporating all inputs (rNE + binned NE + behavior) performed best, achieving a median training correlation of 0.69 and a median held-out correlation of 0.65, compared to 0.55 and 0.53, respectively, for the NE-only baseline model. Adding binned NE or behavioral variables each improved performance incrementally, with the binned NE model showing the most consistent generalization as reflected by the negligible drop between training and held-out correlation (0.62 vs. 0.61). In addition to these promising correlations, RMSE was low and consistent across models (training: 0.02–0.03 ΔF/F; held-out: 0.02–0.04ΔF/F), suggesting that the overall signal magnitude was well-preserved across variants.

**Figure 3.**
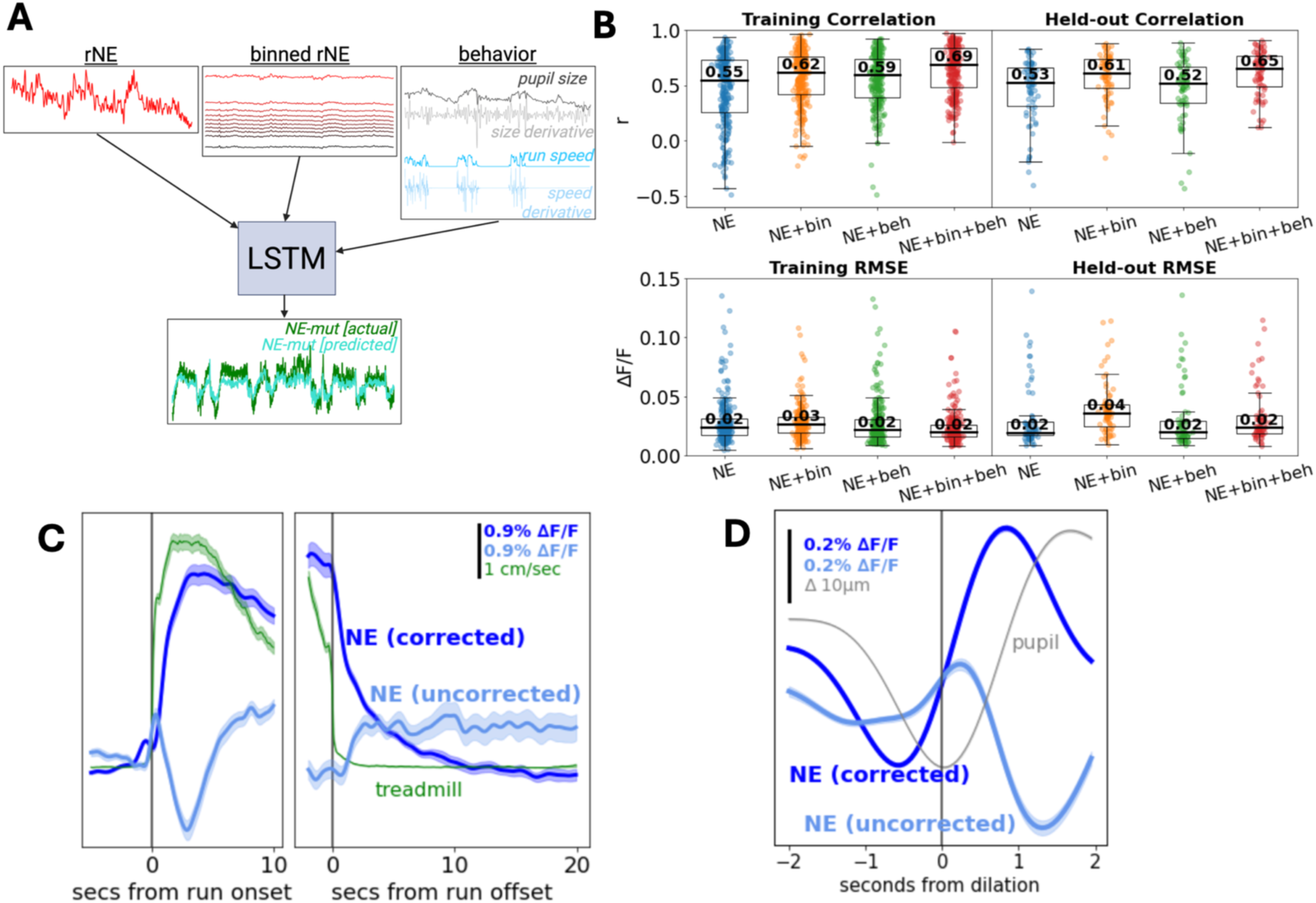
Modeling hemodynamic responses from behavior and NE signals allows post-hoc artifact removal. **A)** Schematic of the LSTM-based modeling approach. Input variables — rNE, binned rNE, and/or behavioral signals (pupil size, treadmill speed, and their temporal derivatives) — are fed into an LSTM network trained to predict the NE-mut fluorescence signal. Four models were evaluated using different input combinations: rNE alone, rNE + binned rNE, rNE + behavior, and all variables combined. **B)** Model performance evaluated by Pearson correlation (top) and RMSE (bottom) for training (left) and held-out (right) data, shown as swarm plots for all four models. Median values are noted in each plot. Models were trained on 267 two-minute segments from 23 recordings and evaluated on 72 held-out segments from 6 recordings. **C)** Mean onset – and offset-triggered uncorrected and model-correct GRAB_NE_ applied to a legacy dataset in which no control channel was recorded (n = 1,010 onsets and 698 offsets from 99 ROIs and 15 animals). **D)** Mean dilation-triggered uncorrected and model-corrected GRAB_NE_ outside of running periods from the same dataset in C (n = 18,317 dilations from 99 ROIs and 15 animals). Shaded regions in C and D indicate ± SEM.

To validate this approach, we applied the full model to a large dataset collected with GRAB_NE_ alone, for which no static control channel was available (the same dataset described in Supplemental Figure 1). We used the full model’s (NE + bins + behavior) predicted hemodynamic signal to correct the raw fluorescence traces and compared run-triggered responses before and after correction (Figure 3C). Following correction, the prominent hemodynamic artifacts were substantially reduced, revealing a cleaner underlying NE signal. Importantly, the corrected signal retained the sustained elevation in NE previously associated with locomotion^12,14,25–27^, indicating that the correction appeared to selectively remove the artifactual component without ablating true NE dynamics.

We further validated the model-based correction by examining pupil dilation-triggered responses during quiescence in the same dataset (Figure 3D). Whereas the uncorrected signal showed dilation-linked fluctuations that were difficult to dissociate from hemodynamic confounds, the corrected signal revealed clear pupil-size-dependent modulation of NE, consistent with the known relationship between pupil diameter and LC or NE activity^12,14,25–32^. Together, these results demonstrate that our predictive model can recover meaningful NE dynamics from recordings that lack a simultaneous control channel.

### Characterizing the relationship between noradrenergic axon activity and extracellular NE dynamics using predictive correction

A key situation in which the dual-channel method cannot be used is experimental designs in which both imaging channels are occupied by signals of interest. As an example of this scenario, we performed simultaneous two-photon imaging of NE axons expressing GCaMP and extracellular NE reported by rNE in either SS/MO cortex or VIS cortex (Figure 4A). Although cortical noradrenergic signaling is often inferred from LC axonal activity^11,25–27,33^, whether and how axonal firing dynamics are reflected in extracellular NE levels has not been directly tested in awake behaving animals. Because both channels carried biological signal, hemodynamic correction relied entirely on the predictive model described above.

**Figure 4.**
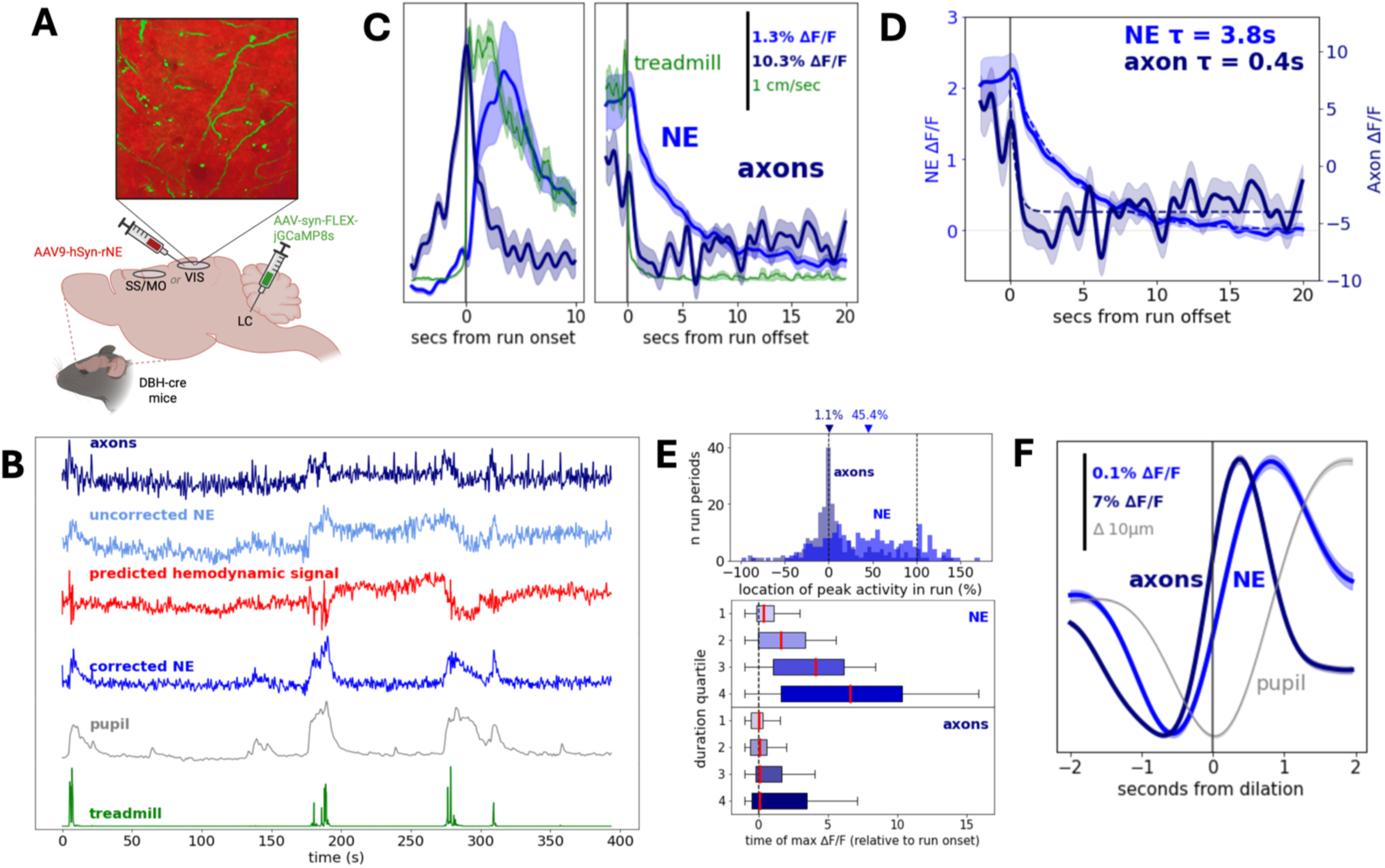
Predictive correction reveals divergence between extracellular NE and LC axon activity during arousal states. **A)** Injection protocol for simultaneous axon-NE imaging experiments. GCaMP8s was expressed in noradrenergic axons by viral injection into the LC of DBH-cre mice. A cranial window was implanted over either the visual or somatosensory/motor cortex and rNE was expressed via viral injection into the cortex. **B)** Example recording traces illustrating the predictive correction. **C)** Mean LC axon activity and model-corrected NE activity aligned to all run onsets and offsets (n=224 onsets and 169 offsets from 35 ROIs and 7 animals). **D)** Exponential decay fits to axonal and NE signals following run offset in C. **E)** (top) Distribution of within-run peak timing for axonal and NE signals, expressed as percent of run duration (0% = onset, 100% = offset; vertical dashed lines). Median values for axons and NE are noted with triangles at the top of the plot (p = 2.59 × 10⁻¹⁵, Kruskal-Wallis test). (bottom) Median peak timing (expressed as seconds after run onset) as a function of run duration quartile. Median values are noted with red lines. **F)** Mean dilation-triggered activity of axon and corrected NE activity outside of running periods (n=4,312 dilations from 38 ROIs and 7 animals). Shaded regions in C, D, and F indicate ± SEM. Quartile bin definitions for run duration and dilation magnitude are provided in Supplemental Fig. 3.

Applying the full predictive model (NE + bins + behavior) to these recordings yielded a corrected NE signal that more closely matched the expected dynamics of extracellular NE. In example recordings, the uncorrected rNE signal appeared noisy and poorly correlated with behavioral state, while the corrected NE signal tracked locomotion and pupil size clearly, with the predicted hemodynamic component accounting for much of the low-frequency structure in the raw trace (Figure 4B).

Both axonal and corrected NE activity were modulated by locomotion, rising at run onset and returning toward baseline following run offset (Figure 4C). However, the timing and dynamics of the two signals differed in ways that suggest extracellular NE is not a trivial readout of axonal firing. To quantify offset dynamics, we restricted analysis to runs without subsequent running in the post-offset window and fit a single exponential decay to each signal. This revealed a slower decay for extracellular NE (τ = 3.8 ± 0.05 sec) than for axonal activity (τ = 0.4 ± 0.05 sec). We initially interpreted this as evidence that extracellular NE remains elevated after axonal activity has subsided (Figure 4D), potentially reflecting uptake and diffusion processes that shape NE availability. However, it is possible this difference in offset timecourse can be mostly explained by differences in the sensor decay kinetics (see below).

Beyond decay dynamics, axonal and extracellular NE signals also differed in when they peaked during a run. Across runs, axonal activity peaked very early (near run onset), with a median peak location of approximately 1.1% of the way through the run (with 0% representing run onset and 100% representing run offset). In contrast, extracellular NE peaked much later, at a median of 45.4% of run duration (Figure 4E, top; axons vs NE p = 2.59 × 10⁻¹⁵, Kruskal-Wallis test). This dissociation was further structured by run duration: binning runs by duration revealed a monotonic relationship in which longer runs were associated with later NE peaks relative to run onset, while axonal activity consistently peaked near the beginning of the run regardless of duration (Figure 4E, bottom). Together, these results suggest that extracellular NE accumulates over the course of a run on a timecourse that is dissociated from instantaneous axonal firing, consistent with prior work demonstrating that LC firing patterns engage distinct NE release dynamics that shape downstream brain state modulation^34^.

Finally, while both axon and NE signals were modulated by pupil dilation outside of running events (Figure 4F), the character of this modulation differed qualitatively: axonal responses appeared more phasic, while extracellular NE and pupil diameter co-varied in a more sustained, sinusoidal manner. This distinction further supports the view that extracellular NE dynamics reflect integrative processes beyond axonal firing alone.

Because axonal GCaMP and extracellular NE were measured with sensors of different kinetics, we asked whether the observed divergence in decay dynamics could be attributed to differences in sensor off-rates rather than true differences in signal timecourse. We deconvolved the NE signal using our previously developed method to remove sensor kinetics^23^ and axonal GCaMP signals using CNMF deconvolution^35^ to remove calcium dynamics (Supplemental Figure 5A). The peak-timing dissociation between axons and NE was preserved following deconvolution (Supplemental Figure 5B; p = 2.33 × 10⁻¹⁵), consistent with the conclusion that this reflects a true biological difference. Furthermore, dilation-triggered responses were qualitatively consistent following deconvolution (Supplemental Figure 5C): axonal activity and extracellular NE rose contemporaneously around pupil dilation events, while extracellular NE showed a broader, more sustained profile than axonal activity.

The decay dynamics of the deconvolved NE signal required careful interpretation (Supplemental Figure 5D). The rNE sensor off-time (∼1.5–3 seconds) is of similar magnitude to the observed raw rNE decay tau (∼3.8 seconds), and the deconvolved trace exhibits ringing artifacts characteristic of Wiener deconvolution under these conditions. Nonetheless, the deconvolved rNE signal decayed near-instantaneously following run offset, suggesting that free extracellular NE is cleared rapidly and that the slow decay observed in the raw signal may be largely attributable to sensor kinetics.

### Development and validation of a behavior-only model for predicting extracellular NE dynamics

We next asked whether NE dynamics could be predicted from behavioral variables alone: an approach that would allow NE to be inferred from any experiment in which locomotion and pupil dilation are tracked. To this end, we trained an LSTM network to predict raw (uncorrected) NE fluorescence from behavior only, using pupil size, run speed, and their derivatives as inputs (Figure 5A). The model was trained on a dataset pooled across legacy GRAB_NE_ recordings and dual-channel recordings (rNE + NE-mut).

**Figure 5.**
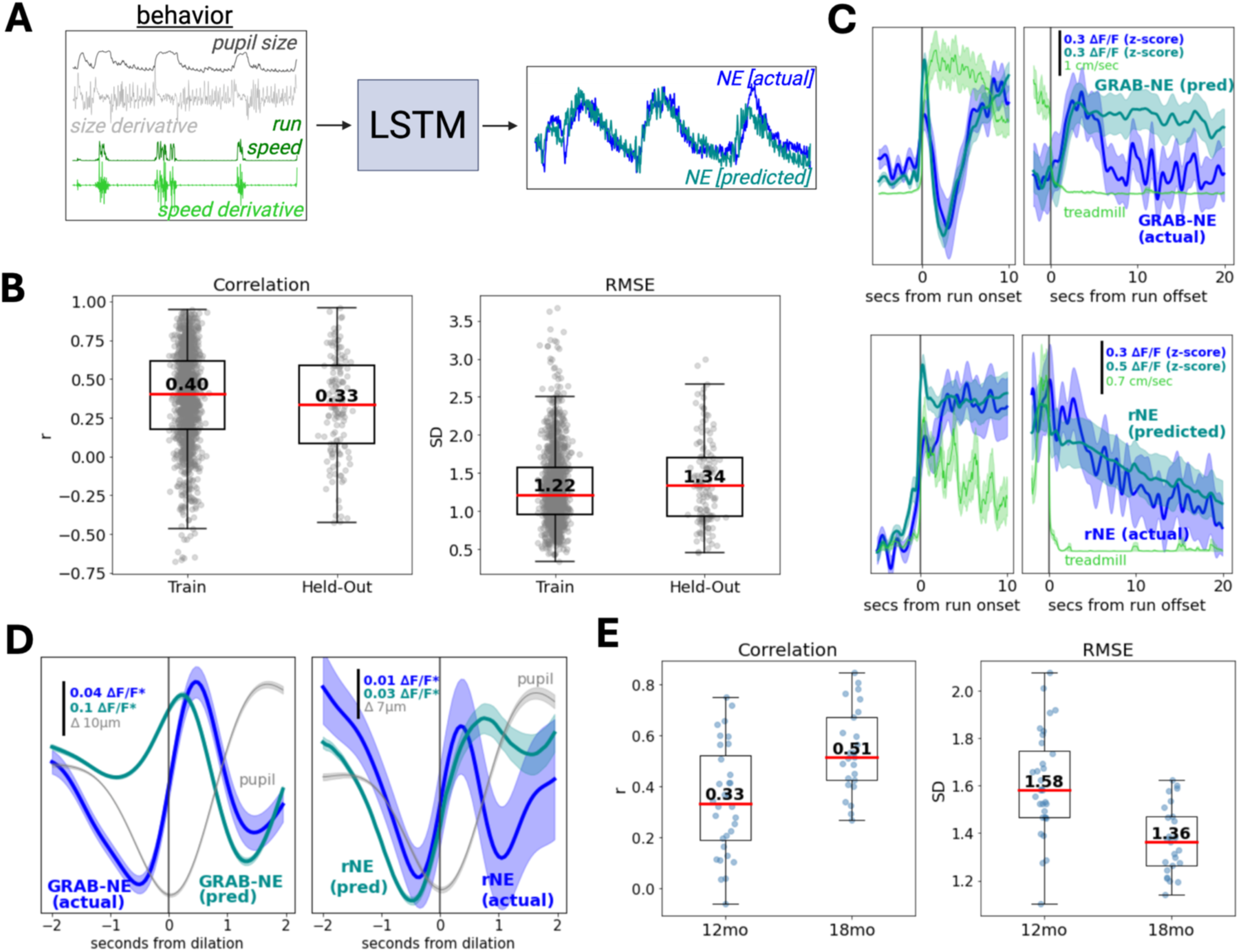
A behavior-only model accurately predicts corrected NE dynamics and generalizes to unseen cohorts. **A)** Schematic of the behavior-only LSTM model. Pupil size, run speed, and their derivatives are used as inputs to predict raw (uncorrected) NE. Data is pooled across legacy and dual-channel recordings. **B)** Model performance evaluated by Pearson correlation (left) and RMSE (right; in standard deviations). Median values are noted in plot. Model was trained on 1,290 2-min segments from 116 recordings and tested on 141 held-out segments from 13 recordings. **C)** Mean run onset- and offset-triggered predicted and recorded GRAB_NE_ (n=54 runs from 11 recordings and 8 animals) and rNE (n=12 runs from 2 recordings and 1 animal) activity from held-out data in B. **D)** Mean dilation onset-triggered predicted and recorded GRAB_NE_ (n=2,269 dilations) and rNE activity (n=444 dilations) from held-out data in B (*ΔF/F z-score). **E)** Model performance on full recordings in an unseen cohort of aged mice. Model is evaluated in 12-month and 20-month cohorts as Pearson correlation (left) and RMSE (right; in standard deviations). Median values are noted in the plot (n=29 12mo and 25 20mo recordings). Shaded regions in C and D indicate ± SEM.

The model achieved fair predictive accuracy on both training and held-out data, with median correlations of 0.40 and 0.33 and RMSE of 1.22 and 1.34 SD respectively (Figure 5B). The similar train and held-out performance suggests that the model generalizes well without overfitting. Mean run-triggered and dilation-triggered activity for held-out GRABNE and rNE scans showed close agreement between predicted and recorded NE (Figures 5C & D). To further validate generalization, we applied the model to an entirely unseen cohort of aged mice (12–20 months). The model performed as well or better on this unseen cohort than even on the training data: median correlations were 0.33 and 0.51 for the 12-month and 20-month cohorts respectively, with RMSE of 1.58 and 1.36 SD (Figure 5E). The good performance on this entirely unseen cohort demonstrates that the model generalizes beyond the training distribution, capturing the behavioral correlates of NE dynamics in animals and recording conditions not represented during training.

While the predictive accuracy of this model was modest, these results demonstrate that behavioral variables alone can indeed recover meaningful features of NE dynamics. Interestingly, an equivalent model trained to predict hemodynamic-corrected NE achieved substantially higher accuracy (Supplementary Figure 6). However, for this prediction, much of the training data was itself model-corrected, and this circularity likely contributes to the inflated performance in predicting corrected NE vs predicting just the raw NE sensor fluorescence.

## DISCUSSION

Here we introduced a tiered framework for inferring more accurate extracellular NE dynamics from fluorescence and behavioral recordings across varying levels of available information. The approaches outlined here should be of general relevance for sensors in which the hemodynamic artifact is of comparable magnitude to the biological signal of interest. The two-channel correction method offers a practical and direct solution to hemodynamic contamination in two-photon NE imaging, requiring no additional light sources or separate control experiments beyond co-expression of an inert fluorescent reporter. We note that no correction method is without limitations: because we found that red fluorophores such as mApple (the fluorophore underlying rNE) are potentially less susceptible to hemodynamic artifacts than green fluorophores (i.e., GFP), correcting rNE with NE-mut may introduce a degree of overcorrection. Conversely, using mApple as a static fluorophore results in systematic under-correction due to its reduced sensitivity to the artifact. A two-FOV approach, in which an NE-sensitive reporter and static fluorophore are expressed in spatially segregated regions and imaged simultaneously, may also be confounded by differences in local vasculature between fields of view. Among the strategies we evaluated, the rNE and NE-mut two-channel approach offered the best practical balance, providing a spatially matched, real-time hemodynamic reference within the same recording session. The regression-based scaling factor (β) estimated during hemodynamic correction partially mitigates the risk for overcorrection inherent to this approach, as it empirically adjusts the contribution of the control channel to best match the sensor signal rather than assuming a fixed 1:1 relationship between the two channels. For recording contexts in which a dedicated reference channel is unavailable, the LSTM-based predictive model offers a complementary solution, enabling post-hoc correction from the recorded NE signal and behavioral variables alone, though model-corrected signals should be treated as improved estimates rather than ground-truth corrections, as prediction quality will degrade in contexts where NE and behavior are weakly coupled or where hemodynamic fluctuations are driven by factors not captured by the input variables. The current model was trained on a relatively limited dataset; nevertheless, training and test performance were strong, and the model successfully removed hemodynamic artifacts in held-out recordings, suggesting the approach is robust. Future studies incorporating larger and more diverse training datasets could further improve generalization across animals, brain regions, and experimental conditions.

Applying these correction methods revealed that extracellular NE is graded with respect to behavioral intensity: NE scaled with both run duration and dilation amplitude, and longer runs produced more sustained NE elevations. In line with previous research suggesting that the magnitude of noradrenergic axon activity scales with pupil dilation^26^, we found that extracellular NE levels themselves also scale with dilation size. Oh the other hand, while locomotion speed has been reported to explain little variance in noradrenergic axon activity^27^, we newly found that run duration seems to be a key determinant of NE magnitude, with longer runs producing progressively larger and more sustained responses. These findings extend prior work demonstrating locomotion- and arousal-coupled NE level increases^8,11,13,31^ by showing that NE also encodes the magnitude and duration of a behavioral state rather than simply its presence or absence, consistent with theories of NE function that emphasize its role in dynamically adjusting cortical gain in proportion to behavioral demands^32^.

Simultaneous imaging of LC axons and extracellular NE provided a rare opportunity to directly compare these signals in the same field of view in vivo: a simultaneous comparison that, to our knowledge, has not previously been made at this spatial and temporal resolution. While noradrenergic axon activity is sometimes used as a proxy for extracellular NE levels, and some studies have reported correlations between the two^11,33^, presynaptic release mechanisms^36^ and cortical state dependence^37^, and diffusion and reuptake dynamics^38^ all have the potential to decouple axonal firing from extracellular NE availability. Our results confirmed that axonal activity and extracellular NE are indeed correlated, but revealed that the relationship is not 1:1 – extracellular NE rises with a small lag relative to axonal activity and peaks much later within a run. Furthermore, NE receptor occupancy outlasts LC axon activity following run offset, even as free extracellular NE appears to be cleared rapidly, suggesting that adrenergic signaling persists beyond the period of active LC firing. The peak-timing dissociation persists following deconvolution, supporting a biological rather than artifactual origin. This stands in contrast to a study suggesting a linear relationship between LC firing and prefrontal NE measured by microdialysis^39^, though the much longer timescales of microdialysis measurements make direct comparison difficult.

The question of whether neuromodulator signaling is “global” or “local” has long been debated. While the LC was historically viewed as a homogeneous nucleus projecting uniformly across cortical areas, more recent anatomical and physiological evidence suggests substantial modularity, with some LC neurons projecting broadly and others targeting discrete cortical regions^40–43^. Furthermore, regional differences in axon density and potentially in reuptake transporter expression have also been found^13,44–46^. Our observation of similar NE increases across VIS, SS, and MO cortex during locomotion, consistent with other reports^9,27^, suggests that large, arousal-driven NE responses may be relatively global in nature. Whether finer-grained heterogeneities may exist, such as the local hotspots of pupil-linked NE activity reported elsewhere^14^, remains an open question that our study was not designed to resolve, as we did not differentiate between subregions within each field of view.

Finally, the behavior-only model demonstrates that key features of raw NE dynamics can be recovered from locomotion and pupil signals alone. Despite modest overall predictive accuracy, the model was able to generalize well to an unseen aged cohort. This provides a tool for estimating neuromodulatory state in experiments where fluorescence recording is unavailable, and offers a behavioral baseline against which richer recordings can be compared. A key limitation is that the model captures only variance in NE that covaries with the behavioral inputs; fluctuations driven by sensory stimuli, internal states, or other factors not reflected in locomotion and pupil size will not be recovered. Future extensions incorporating additional behavioral or physiological variables may improve predictive power and broaden applicability.

Together, these correction and modeling approaches form a practical toolkit for isolating true NE dynamics across varying levels of recording information. By making robust NE measurement accessible across a broader range of experimental contexts, this framework enables more reliable interpretation of the growing body of genetically encoded NE sensor data, and reveals that the spatiotemporal structure of cortical NE signaling is richer and more graded than would be apparent from uncorrected recordings.

## METHODS

### Animals, surgery, and injection of viral vectors

All procedures were carried out in accordance with the ethical guidelines of the National Institutes of Health and were approved by the Institutional Animal Care and Use Committee (IACUC) of Baylor College of Medicine. This study used a total of 49 mice aged 2 to 20 months (n=27 male, n=22 female).

All surgeries were performed under isoflurane anesthesia. Wild-type C57Bl/6 mice (or negative genotype littermates on a C57Bl/6 background) were used unless otherwise noted. The head was shaved over the dorsal portion of the skull and bupivacaine 0.25% (0.1 ml) and ketoprofen (5 mg/kg) were administered subcutaneously. The skin was pulled back and glued in place using Vetbond (3M). A headbar (steel washer with outer diameter 0.438”, inner diameter 0.314”, thickness 0.016”) was affixed to the skull using dental cement (C&B Metabond). A micromotor rotary drill (Foredom) fitted with a 0.6 mm tungsten carbide bur was used to thin and remove the skull over the target area. Viral injections were made using a borosilicate glass pipette pulled to a 40 µm tip, attached to a syringe and Micro4 MicroSyringe pump controller. The dura was removed and a #1 borosilicate glass coverslip (0.15 ± 0.02 mm) was glued in place with Loctite. Animals recovered on a heated disk until sternal recumbency was regained and were monitored for 3 days post-surgery. Stereotactic coordinates for window placement were as follows: VIS: 2.8 mm lateral, 1 mm anterior lambda; VIS/SS: 2.8 mm lateral, 2.25 mm posterior bregma, SS: 3.25 mm lateral, 0.3 mm posterior bregma, MO: 1.5 mm lateral, 1.5 mm anterior bregma, SS/MO: 2.25 mm lateral, 0.5 mm anterior bregma.

### Dual-channel static fluorophore recordings (rNE + NE-mut)

Mice were implanted with a 4 mm cranial window over VIS (n=2), a 5mm window over SS/VIS (n=2) or a 3mm window over MO (n=3).

### Dual-channel static fluorophore recordings (GRABNE + mApple)

Mice were implanted with a 4 mm cranial window over VIS (n=1), SS (n=1), or MO (n=1) and injected with 1300nl of a 4:1 mix of AAV9-hSyn-mApple and AAV-hSyn-NE2h at a depth of 200-700µm.

### Dual-field recordings

2 mice were implanted with a 4mm cranial window over VIS. Small amounts of AAV-hSyn-NE2h and AAV-hSyn-NEmut were injected in spatially segregated locations within the window (250nl each, at a depth of 300µm).

### Single-channel and GFP control recordings

Mice received a 4mm cranial window over VIS and were injected with either 500-2000nl of AAV-hSyn-NE2h (n=16) or a GFP-only construct (n=1) at a depth of 200-700µm.

### Axon/rNE recordings

DBH-Cre mice first received an LC injection either under ketamine/xylazine or isoflurane anesthesia. A burr hole was drilled and a micropipette was positioned at AP: -5.45, ML: 1.2, DV: -3.9; 1600-2000 nl of AAV-syn-FLEX-jGCaMP8s was injected over 160 seconds. The skin was sutured with braided absorbable sutures (McKesson) and the animal was allowed to recover for at least 1 week. A headbar and 4mm cranial window was then implanted over VIS or SS/MO using the procedures above, into which 1000-2000nl of AV9-hSyn-rNE0.5 (n=3) or AAV9-hSyn-rNE1.0 (n=4) was injected at a depth of 300µm.

### Aged mouse recordings

13 mice were implanted with a 4mm cranial window over SS/MO. AAV9-hSyn-rNE1.0 and AAV-hSyn-ACh4.3 were injected in spatially segregated locations within the window (250nl each, at a depth of 300µm).

#### Locomotion and pupillometry

Mice were head-fixed on a cylindrical treadmill, and locomotion was tracked using an optical encoder. Running was defined as speeds ≥1 cm/sec sustained for ≥1 sec. Pupil images were captured at 20 fps and analyzed using DeepLabCut to extract pupil radius (converted to mm using a conversion factor based on corner-to-corner eye distance). Dilation and constriction events were defined as periods of increasing or decreasing pupil radius lasting >1 sec with average speed >0.02 mm/sec. Full methodological details have been described previously^23^.

#### Imaging

All scans were recorded using a two-photon fast resonant scanning system (ThorLabs). Most scans were acquired at ∼30 fps (n=268 scans), with a small subset acquired at other frame rates (n=33, 5.7-58.3 fps). Excitation wavelengths were either 980-1000nm (simultaneous axon GCaMP and rNE scans), 1020-1055 (rNE and NE-mut or mApple and GRAB_NE_), 920nm (GRAB_NE_ or GFP-only scans), or 980-1020nm (“aged” mice rNE and GRAB_ACh_ scans). Objectives ranged from 10-25x (Nikon). Output power was kept around 15-40mW. The green emission filter was 520-550nm and the red emission filter was 620-670nm. Most scans were imaged with an average FOV size around 200×200µm with the exception of scans the aged mice cohort, which were imaged at an average FOV size of around 1,700×1,700µm.

#### Motion correction and selection of FOVs

All scans were corrected for motion in X and Y using a two-step phase correlation algorithm (large-scale followed by local, windowed correction). Scans with poor motion correction (RMS shift >4 µm). Full details of the motion correction have been described previously^23^.

Fluorescence traces for all scans were determined by taking the average fluorescence over time of all pixels within a field-of-view mask (“FOV mask”). Unless otherwise noted, masks were defined using pixels with average brightness values between the 99.9th percentile/2 and the 99.9th percentile after removing scan borders, then eroded using a 7×7-pixel kernel. Exceptions and additional details are described below.

For scans in which only a single channel was recorded (legacy GRAB_NE_ scans, two-field static fluorophore scans, and GFP scans), masks were determined using the brightness criteria above.

For dual-channel scans, the mask was first determined on the green channel using the brightness criteria above. This same mask was then applied to the red channel, as the green channel was selected as the reference due to its relative brightness.

For axon scans, the green channel (GCaMP) mask was determined by calculating the overlap between a manually drawn mask over visible axons and a brightness mask (pixels with average brightness >99th percentile/2 after removing borders). The red channel (rNE) mask was determined using the standard brightness criteria above (but instead bounded at the lower end by the 99th percentile/2).

For scans in the aged mice cohorts, rNE1.0 and GRAB_ACh_3.0^22^ were expressed and recorded simultaneously in spatially segregated regions of the cranial window. Red and green channel masks were determined by calculating the overlap between manually drawn masks over the respective sensor areas, and brightness masks (green channel: >99.9th percentile/2; red channel: >99th percentile/2), after removing scan borders and eroding with a 7×7-pixel kernel. Only data from the rNE masks were used in this study.

Binned masks for hemodynamic modeling were determined by computing the average fluorescence over time across 10 equal bins spanning the darkest (0–10%) to brightest (90–100%) pixels after removing border pixels. For axon scans, pixels within the axon mask described above were excluded from the bins to avoid potential bleedthrough from the green (GCaMP) channel. For aged mice cohort scans, bins were computed only within the manually drawn rNE region.

#### Fluorescence trace preprocessing and scan exclusion criteria

Unless otherwise noted, all fluorescence traces were preprocessed identically prior to any analysis or visualization. The first 20 seconds were discarded to allow imaging stabilization, traces were detrended (4th-order polynomial) to correct for photobleaching, low-pass filtered at 1 Hz, and converted to ΔF/F using a rolling 5th-percentile baseline (20-minute non-causal window). Several exclusion criteria were applied: traces with negative baseline values were discarded to avoid ΔF/F normalization artifacts; scans with abnormally high ΔF/F range relative to baseline (ratio of fluorescence range to 5th-percentile baseline greater than 5) were discarded as outliers; dual-channel static fluorophore recordings in which the sensor and control channels showed substantially similar deflection patterns were excluded due to suspected channel cross-contamination/ bleedthrough; and scans containing large unexplained jumps in fluorescence (at least 20x the average derivative) were excluded.

#### Peri-dilation, peri-run, and peri-spike traces

Run onset and offset, and pupil dilation onsets were defined as described above (“Locomotion and pupillometry”). For all peri-event traces (peri-dilation, peri-run onset, and peri-run offset), the baseline was defined as the mean of the full pre-event window, except for axon peri-run traces. Because axon activity begins ramping prior to run onset, making the immediate pre-onset period unsuitable as a baseline, axon peri-run traces were baselined to the first 3 seconds of the 5-second pre-onset window, excluding the 2 seconds immediately preceding run onset. For peri-offset traces, both signals were baselined to the same pre-onset period from the corresponding run onset. Variability is reported as standard error of the mean.

#### Correction of hemodynamic artifact

Sensor (GRAB_NE_ or rNE) and control (either recorded or predicted NE-mut) signals were preprocessed as described above. Hemodynamic correction was then applied as follows:

A scaling factor (β) was estimated via linear regression of the control signal onto the sensor signal using only quiescent periods (i.e. excluding running periods and a 5-second pre-onset and 15-second post-offset buffer). The mean offset between the quiescent sensor and control signals was then used to align the control signal to the sensor baseline. The hemodynamic correction was then applied to the full trace as:

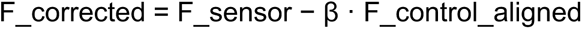

The corrected trace was then baseline-shifted by the median of the quiescent sensor frames and lowpass filtered to attenuate high-frequency noise introduced by the subtraction procedure.

#### Deconvolution

The full deconvolution method is described in detail previously^23^. Briefly, fluorescence signals were preprocessed as described above and then deconvolved using Wiener deconvolution, which models the fluorescence trace as the convolution of the underlying neurotransmitter concentration with a single falling-exponential impulse response function characterizing the sensor kinetics. The regularization parameter λ was selected empirically on a per-scan basis. τ_off_ values used were 1.5 s (rNE0.5; VIS scans) and 3.1 s (rNE1.0; SS/MO scans).

Axon fluorescence, extracted from the axon mask described above, was deconvolved using constrained non-negative matrix factorization (CNMF)^35^. Deconvolved axon traces were subsequently lowpass filtered at 1 Hz to remove high-frequency noise for visualization purposes.

#### Hemodynamic predictive modelling

Fluorescence traces (rNE and NE-mut) were preprocessed as described above (detrending, 1 Hz lowpass filter, ΔF/F normalization). rNE data were binned into 10 fluorescence percentile bins as described above (“Motion correction and selection of FOVs”), preprocessed identically, and jointly z-scored across bins. Behavioral covariates included pupil radius, (1 Hz lowpass filtered) locomotion speed (median filtered), and their first derivatives. All fluorescence and behavior signals were interpolated to a common timebase (20 fps). Continuous recordings were segmented into non-overlapping 2-minute windows. Only recordings with concurrent pupil data were included so that all model variants could be compared on an identical dataset. Data were pooled across VIS, SS, and MO.

We trained an LSTM-based regressor (single LSTM layer, 256 hidden units; fully connected layer, 256 units, ReLU; dropout p = 0.1; linear output) to predict mutant sensor fluorescence from four input combinations: (1) rNE alone, (2) rNE + brightness bins, (3) rNE + behavioral covariates, and (4) rNE + brightness bins + behavioral covariates. Models were trained using MSE loss and the Adam optimizer (lr = 1×10⁻³, 50 epochs, batch size 32). Train/test splits were performed at the level of whole recordings (∼20% held out per region; V1: 1, S1: 2, M1: 3). Prediction accuracy was quantified using Pearson correlation and root mean squared error (RMSE) on held-out data pooled across regions.

#### Behavioral to NE predictive modelling

NE fluorescence traces were preprocessed as described above (detrending, 1 Hz lowpass filter, ΔF/F normalization) and z-scored across the full recording. Behavioral covariates included pupil radius, locomotion speed (1 Hz lowpass filtered), and their first derivatives, each z-scored across the full recording. All signals were interpolated to a common timebase (20 fps). Continuous recordings were segmented into non-overlapping 2-minute windows. Data were pooled across recordings from dual-channel recordings (rNE + NE-mut) as well as legacy recordings (GRAB_NE_ only). Sensor type (rNE vs. GRAB_NE_) was included as a one-hot encoded covariate.

We trained an LSTM-based regressor (identical architecture to above) to predict z-scored (raw/uncorrected) NE fluorescence from the behavioral covariates. The model was trained using a Pearson correlation loss and the Adam optimizer (lr = 1×10⁻³, 50 epochs, batch size 32). Train/test splits were performed at the level of whole recordings (∼10% held out, randomly sampled without stratification). Prediction accuracy was quantified using Pearson correlation and RMSE on held-out data.

A second model was trained using an identical architecture and training procedure to predict hemodynamic-corrected NE fluorescence rather than raw NE fluorescence. Traces were corrected as described above prior to modeling, using either a recorded or predicted hemodynamic signal (NE-mut). Behavioral covariates included pupil radius, pupil derivative, and locomotion speed only (no locomotion acceleration or sensor one-hot encoding). Training data were pooled across dual-channel recordings, legacy single-channel recordings, and axon + rNE recordings.

## ACKNOWLEDGEMENTS

We would like to thank the Jeannie Chin laboratory for providing mice used in the “aged mouse” and the “axon/rNE” cohorts used in this study. We would also like to thank the Andreas Tolias lab for providing resources and equipment. J.R. was supported by NIH awards R01 NS128901 and R34 NS137454. E.N. was supported by NIH NRSA Fellowship F31NS122428. B.R.M was supported by a Dementia Australia Research Foundation Project Grant and the Australian Research Council (DECRA DE260101935). Y.L. was supported by the National Key R&D Program of China (2021YFF0502904) and the National Natural Science Foundation of China (32525003). J.M.S. was supported by the National Health and Medical Research Council (GNT1193857) and Australian Research Council (DP240101295 and DP250102186). Some figures were created with the assistance of Biorender.

## AUTHOR CONTRIBUTIONS

Conceptualization, E.N., J.R.; Methodology, E.N., B.R.M., J.R.; Formal Analysis, E.N.; Investigation, E.N., N.Z., P.Y., J.F.; Resources, Y.L., J.M.S., J.R.; Data Curation, E.N.; Writing – Original Draft, E.N.; Writing – Review & Editing, B.R.M, Y.L., J.M.S., J.R.; Visualization, E.N.; Supervision, Y.L., J.M.S., J.R.; Funding Acquisition, E.N., B.R.M., Y.L., J.M.S., J.R.

---

**Supplemental Figure 1.**
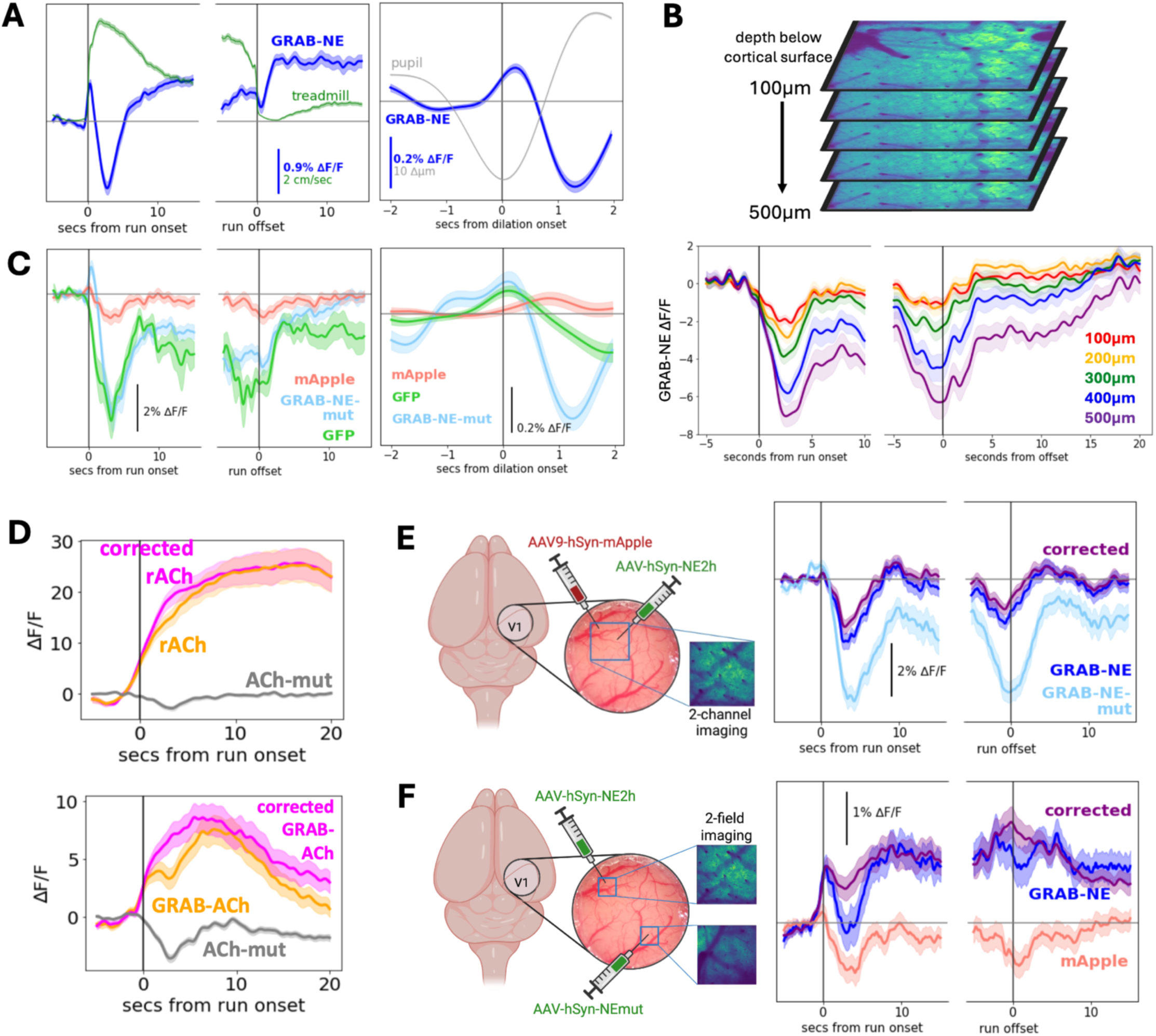
**A)** GRAB_NE_ fluorescence aligned to all run onsets and offsets (left; n=1,414 run periods from 120 ROIs and 18 mice), or to all dilation onsets during quiescence (right; n=18,382 dilation periods from 101 ROIs). Shading is +/- standard error. **B)** Recording five simultaneous planes of GRAB_NE_ in the z-direction at 100µm intervals. Aligning to run onset and run offset (n=44 run periods) reveals a stronger artifact at deeper depths. **C)** Activity traces of various control indicators aligned to run onset and offset (left; mApple n=108 runs, GRAB-NE-mut n=418, GFP n=15), or to dilation onset during quiescence (right; mApple n=1,517 dilations, NE-mut n=6,267, GFP n=801). z (top) rACh and ACh-mut recorded simultaneously in a two-channel paradigm compared with corrected rACh aligned to run onset; (bottom) GRAB_ACh_ and ACh-mut recorded simultaneously in a two-field paradigm compared with corrected GRAB_ACh_ aligned to run onset. [reproduced from Neyhart et al., 2024^23^] **E)** (left) Schematic of two-channel recording method with overlapping expression of GRAB_NE_ and mApple. (right) GRAB_NE_, mApple, and corrected GRAB_NE_ traces aligned to run onset and offset (n=141 run periods from 14 ROIs). **F)** (left) Schematic of two-field recording method with non-overlapping expression of GRAB_NE_ and NE-mut in spatially segregated areas of the cranial window. (right) GRAB_NE_, NE-mut, and corrected GRAB_NE_ traces aligned to run onset and offset (n=71 runs from 6 ROIs).

**Supplemental Figure 2.**
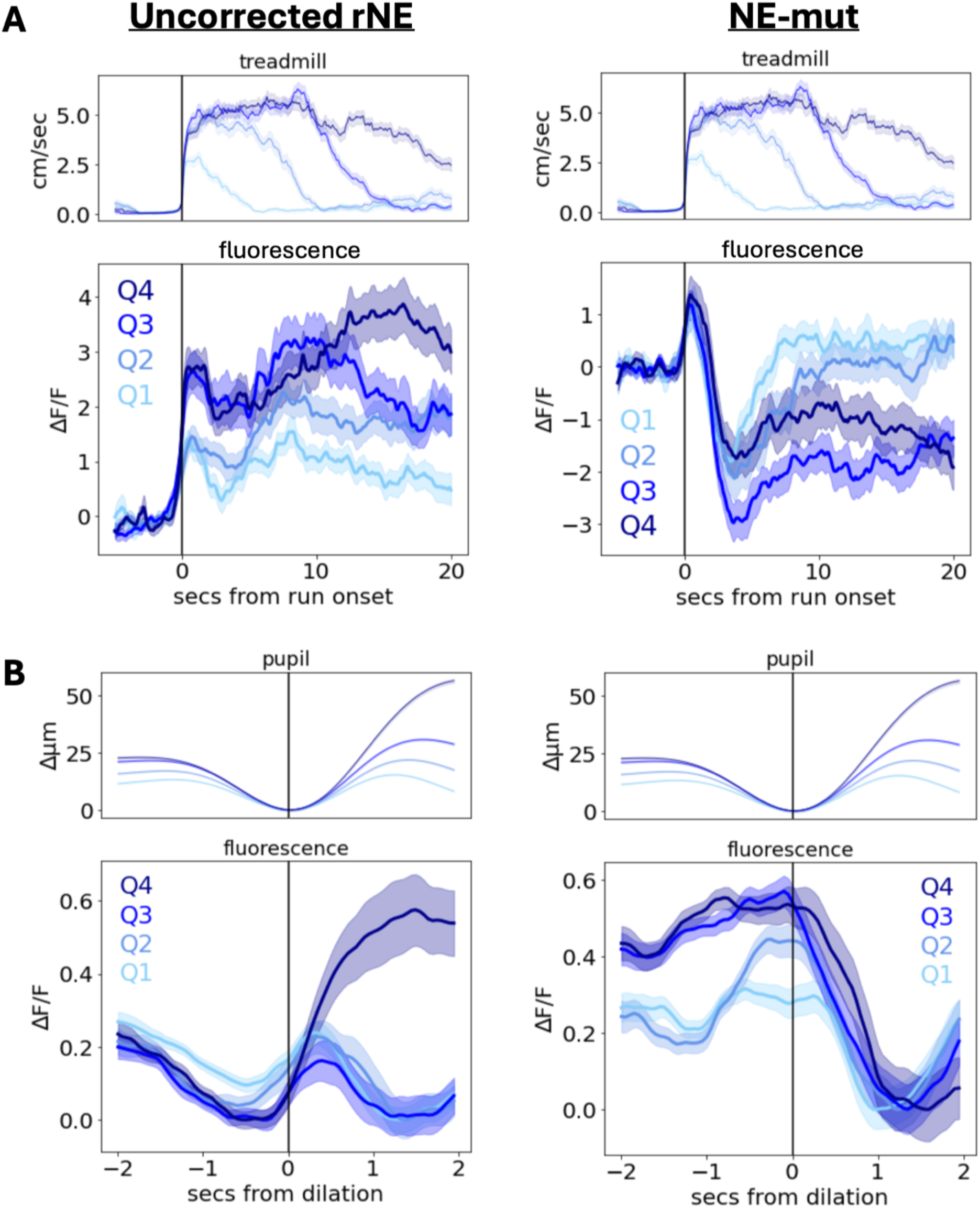
**A)** All run onsets sorted by run duration: (left) uncorrected rNE, (right) NE-mut. **B)** All pupil dilations sorted by dilation amplitude: (left) uncorrected rNE, (right) NE-mut).

**Supplemental Figure 3.**
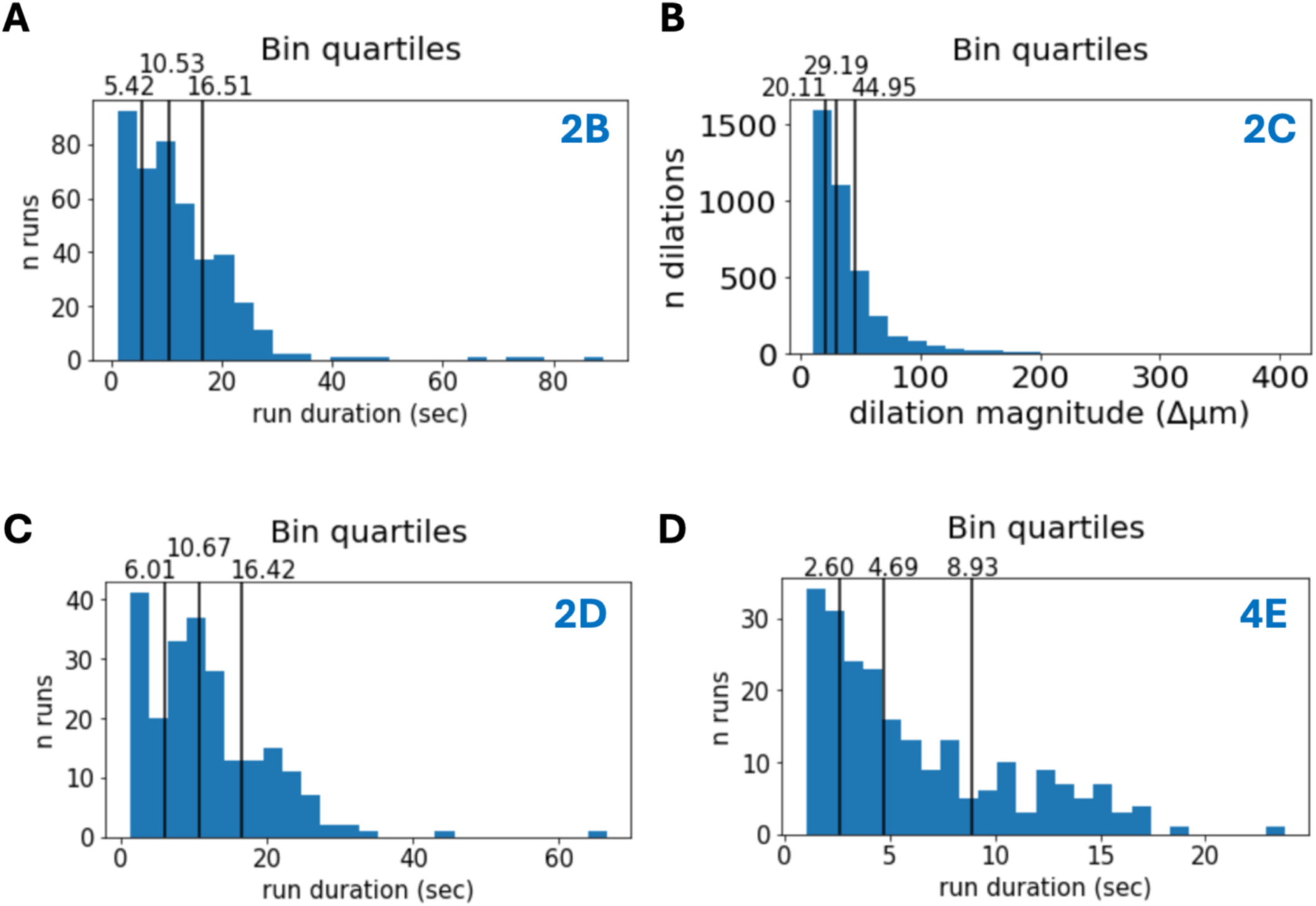
Bin quartiles for various plots. Main figure referenced in top right corner of each panel.

**Supplemental Figure 4.**
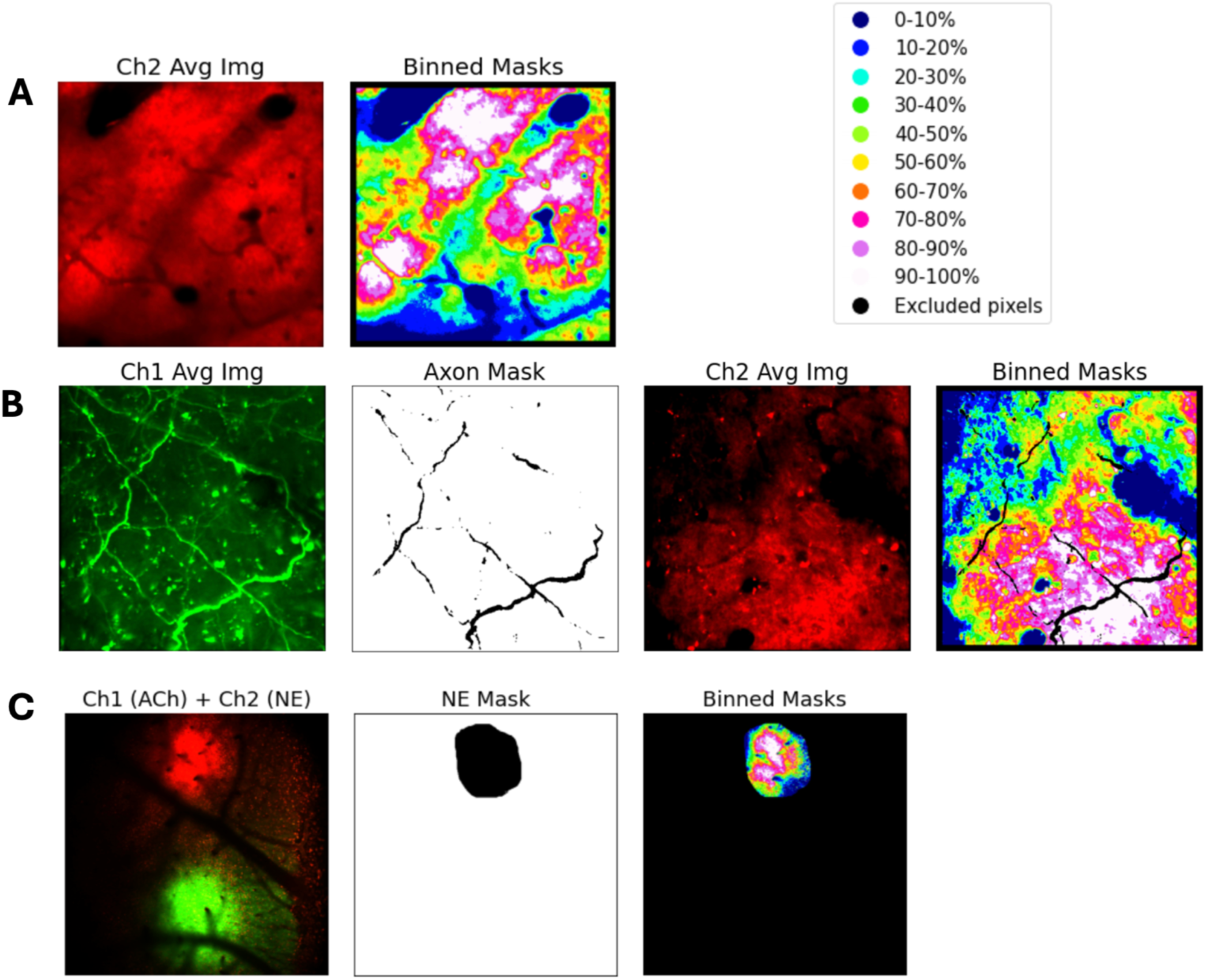
Binned masks used in hemodynamic prediction. **A)** Intensity-based spatial bins computed across the full field of view for legacy and dual-channel static fluorophore scans. Pixels were ranked by average intensity in the red channel and divided into 10 equal bins spanning the darkest (0–10%) to brightest (90–100%) pixels; border pixels were excluded from all bins. **B)** Bins computed for dual axon + rNE recordings. Axon mask pixels were excluded prior to computing average red channel intensity. **C)** Bins computed for scans containing multiple spatially segregated sensors. Bins were determined exclusively within a manually drawn region of interest delineating the rNE expression area (”NE mask”), rather than across the full scan area.

**Supplemental Figure 5.**
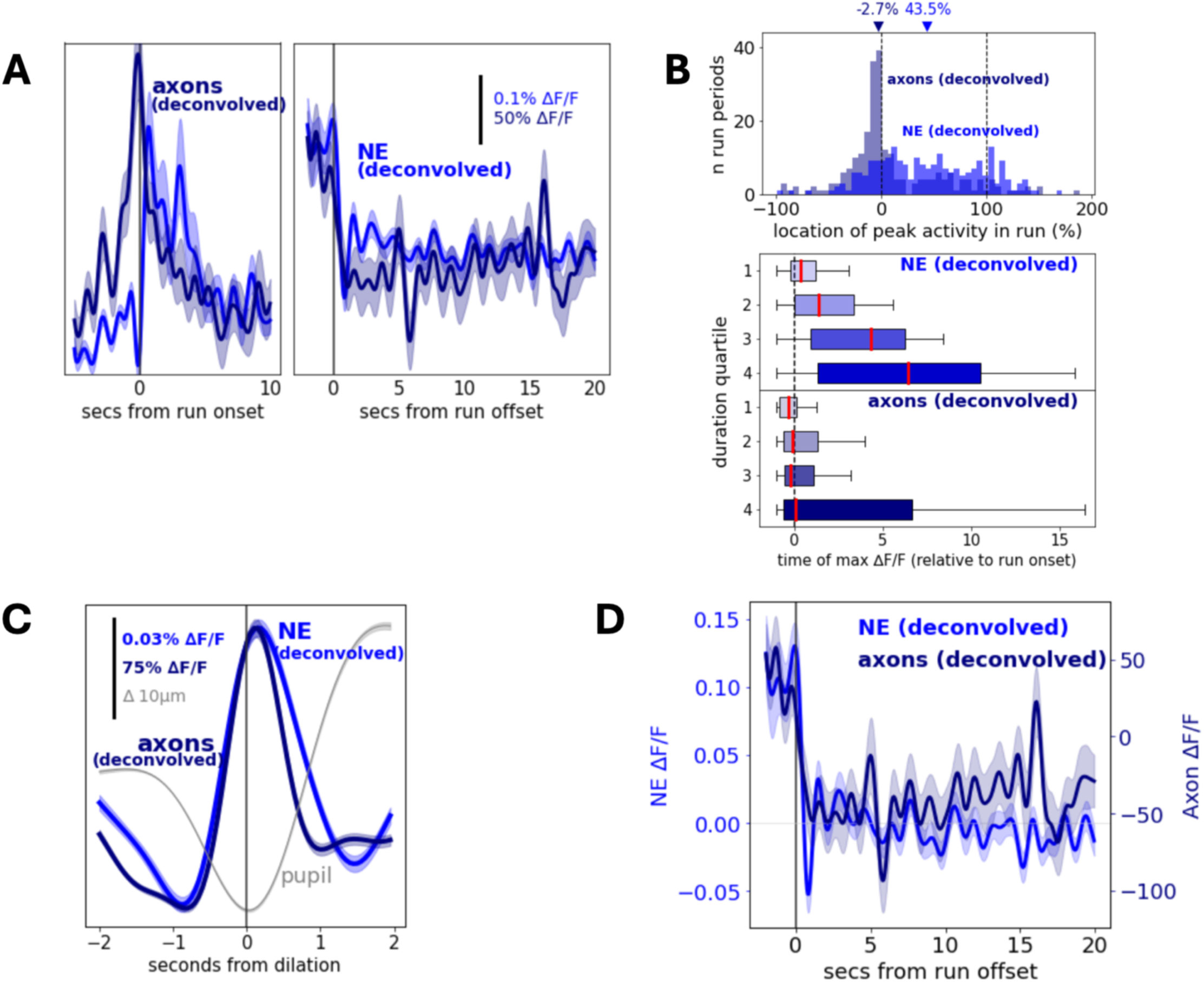
Results from Figure 4, instead with CNMF-deconvolved LC axon activity and sensor-deconvolved corrected NE. **A)** Mean LC axon activity and model-corrected NE activity aligned to all run onsets and offsets. **B)** (top) Distribution of within-run peak timing for axonal and NE signals, expressed as percent of run duration (p = 2.33 × 10⁻¹⁵, Kruskal-Wallis test). (bottom) Median peak timing (expressed as seconds after run onset) as a function of run duration quartile. **C)** Mean dilation-triggered activity of axon and corrected NE activity outside of running periods. **D)** Mean run offset-triggered activity in A, replicated for clarity.

**Supplemental Figure 6.**
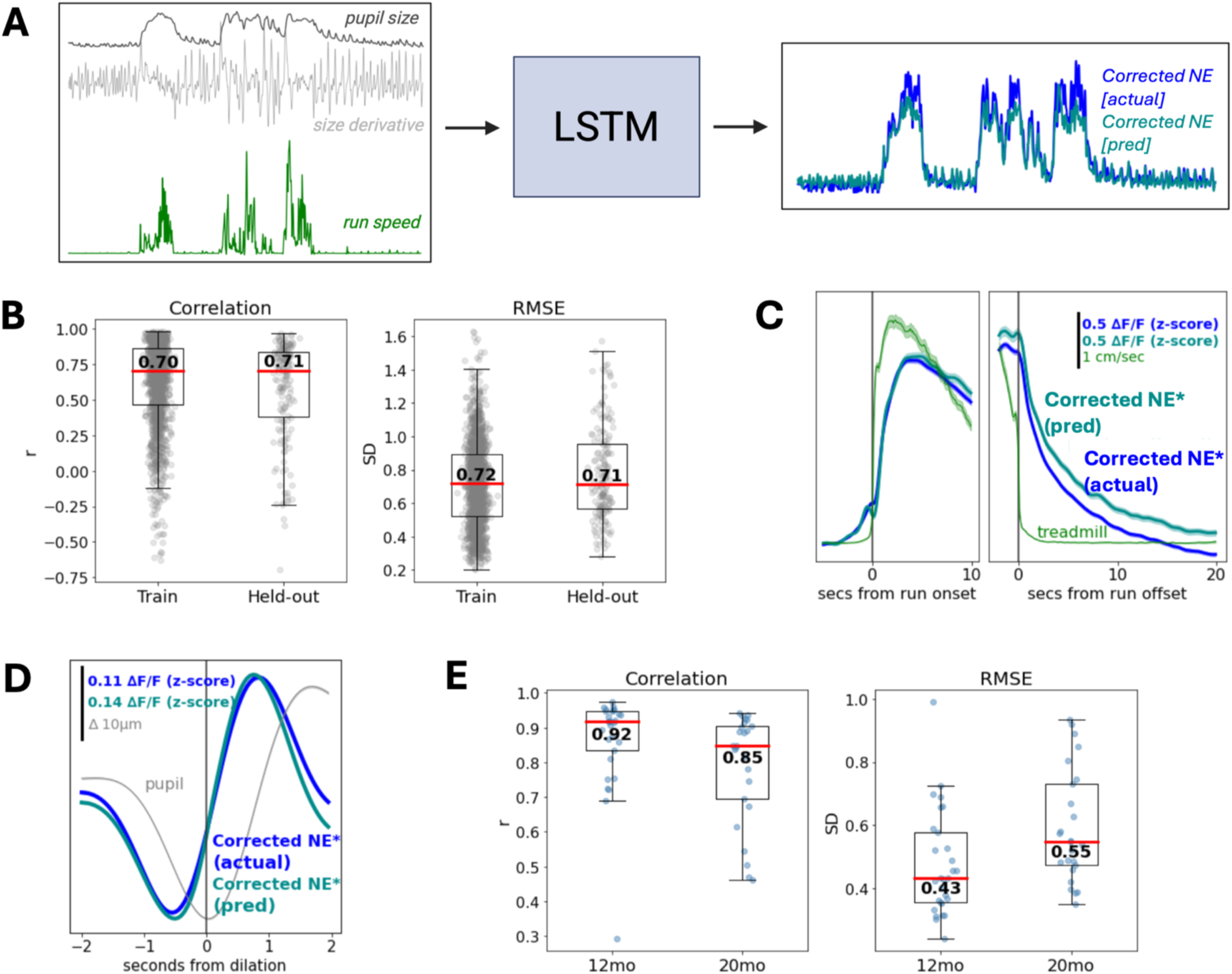
A behavior-only model accurately predicts corrected NE dynamics and generalizes to unseen cohorts. **A**) Schematic of the behavior-only LSTM model. Pupil size, its derivative, and run speed are used as inputs to predict hemodynamic-corrected NE. pooled across legacy, dual-channel, and axon imaging recordings, with hemodynamic correction performed using either recorded or predicted hemodynamic signals depending on data availability. **B)** Model performance evaluated by Pearson correlation (left) and RMSE (right; in standard deviations). Median values are noted in plot. Model was trained on 1,605 two-minute segments from 146 scans and 27 animals and 171 segments from 16 scans. **C)** Mean run onset- and offset-triggered predicted and recorded (corrected) NE activity (*data combined across GRAB_NE_ and rNE recordings) from all training and test data in B (n=1,080 run periods). **D)** Mean dilation-triggered activity of predicted and recorded (corrected) NE activity (*data combined across GRAB_NE_ and rNE recordings) outside of running periods in the same dataset in B and C (n=27,093 dilations). **E)** Model performance on full recordings from an unseen cohort of aged mice. Model is evaluated in 12-month and 20-month cohorts as Pearson correlation (left) and RMSE (right; in standard deviations). Median values are noted in the plot (n=29 12mo and 25 20mo recordings). Shaded regions in C and D indicate ± SEM.

## Notes

### Competing Interest Statement

The authors have declared no competing interest.

### Summary of Updates

Added Author Contributions, an expanded Acknowledgements section, and another author

